# A comprehensive data repository of environmental toxicants exposed mouse epigenomes from the TaRGET II Consortium

**DOI:** 10.1101/2025.07.28.667245

**Authors:** Prashant K. Kuntala, Benpeng Miao, Deepak Purushotam, Bo Zhang, Daofeng Li, Ting Wang

**Affiliations:** The Edison Family Center for Genome Sciences and Systems Biology, Washington University School of Medicine, St. Louis, MO; McDonnell Genome Institute, Washington University School of Medicine, St. Louis, MO, USA; Department of Genetics, Washington University School of Medicine, St. Louis, MO; Center of Regenerative Medicine, Washington University School of Medicine, St. Louis, MO; Department of Developmental Biology, Washington University School of Medicine, St. Louis, MO

## Abstract

The Toxicant Exposures and Responses by Genomic and Epigenomic Regulators of Transcription (TaRGET) is a multiphase program that aims to understand how environmental factors contribute to disease susceptibility using toxicant-exposed mouse models. Here, we introduce the TaRGET II Data Portal (https://data.targetepigenomics.org/), a repository of exposomes from mice exposed to environmental toxicants, including arsenic (As), lead (Pb), bisphenol A (BPA), tributyltin (TBT), di(2-ethylhexyl) phthalate (DEHP), tetrachlorodibenzo-p-dioxin (TCDD), and air pollution (PM2.5). Sequencing assays capturing changes in chromatin accessibility, DNA methylation, gene expression, and post-translational histone modifications from multiple centers were quality-controlled and uniformly processed. The datasets cover multiple tissues collected at four time points: 3 weeks, 5 weeks, 20 weeks, and 40 weeks. The TaRGET II Data Portal offers an efficient way to browse, search, visualize, and download relevant datasets and associated metadata, serving as a key resource for studying the impact of environmental toxicant exposures on disease susceptibility for the broader scientific community.

## Introduction

Understanding both the immediate and long-term effects of environmental toxicant exposure on the epigenome in various tissues is crucial for developing effective intervention strategies and accurate disease prognosis. However, the impact of dosage, duration, and timing of exposure on regulatory pathways that contribute to adverse health outcomes remains unclear. Furthermore, evaluating the feasibility of using surrogate tissues, such as blood and skin, as alternatives to more challenging-to-obtain target tissues like the heart and liver is vital for diagnosis and for accurately assessing the extent of toxicant exposure ^1–7^.

The Toxicant Exposures and Responses by Genomic and Epigenomic Regulators of Transcription (TaRGET)^7^ consortium is a multiphase program led by the National Institute of Environmental Health Sciences (NIEHS) to investigate how environmental factors contribute to disease susceptibility through changes in the epigenome. The second phase, TaRGET II, focuses on exploring the conservation of the epigenetic landscape across various tissues and cell types in toxicant-exposed mouse models, with the intent to improve the design and analysis of human studies where such tissues are difficult to obtain.

The consortium comprises five data production centers and a data coordination center (DCC). The production centers generated bio-samples by exposing mice to various environmental toxicants and collecting both target and surrogate tissues over time. They conducted biochemical assays to measure changes in chromatin accessibility, DNA methylation, gene expression, and both active and repressive histone post-translational modifications (hPTMs). The DCC developed quality control and analysis pipelines for these assays, curated metadata, created a unified sample submission process, and developed resources for disseminating raw and processed data, along with preliminary analysis results.

Here, we introduce the TaRGET II Data Portal, a comprehensive resource of next-generation sequencing (NGS) datasets that aids in understanding the role of environmental toxicants in disease susceptibility. We highlight available datasets and demonstrate how users can browse, search, download, and visualize data using the WashU Epigenome Browser, showcasing its value to the broader scientific community.

## Results

The TaRGET DCC developed a web-based data portal to host metadata and quality-control metrics for all data generated by the TaRGET II consortium. Built on the latest cloud-compliant technologies using Amazon Web Services (AWS), the portal is designed to scale across various project types and use cases. In the TaRGET II data portal, the top-level data model is an experiment, which uniquely represents a biochemical assay conducted under defined conditions, including exposure, tissue type, age, and sex. Data production centers generated multiple biological replicates per experiment, and currently, 3,091 experiments are available on the portal.

### Exposome Profiling

The genomic datasets include several key assays: ATAC-seq^8^ for assessing chromatin accessibility, RNA-seq^9^ for measuring gene expression, and both Whole Genome Bisulfite Sequencing (WGBS)^10^ and Reduced Representation Bisulfite Sequencing (RRBS)^11^ for profiling DNA methylation. In some cases, Illumina BeadChip arrays^12^ were used to assess 5-methyl-cytosine (5mC) and 5-hydroxy-methyl-cytosine (5hmC). Histone modifications were profiled via ChIP-seq^12^ for markers such as H3K27ac, H3K4me1, H3K4me3, H3K27me3, and H3K9me3. Methylation BeadChip arrays^12^ were primarily performed on blood and cortex samples, while ChIP-seq for active and repressive histone modifications was mainly conducted on liver samples. All genomic assays available through the TaRGET II data portal and accessible to the public are summarized in **Table 1**.

**Table 1.** Total experiments (N=3091) summarized by Exposure, Tissue and Age. Exposures include: Arsenic (As), lead (PB), bisphenol A (BPA), tributyltin (TBT), di(2-ethylhexyl) phthalate (DEHP), tetrachlorodibenzo-p-dioxin (TCDD), and air pollution or Particulate Matter (PM2.5). ^*^ PM2.5 (JHU), PM2.5 (CHI) are used to differentiate between data production centers that used different exposure paradigms.

|  |  |  |
| --- | --- | --- |
| **Exposure** | Arsenic | 95 |
|  | BPA (10mg) | 193 |
|  | BPA (10ug) | 189 |
|  | DEHP | 210 |
|  | PM2.5 (JHU)* | 170 |
|  | PM2.5 (CHI)* | 219 |
|  | PB | 220 |
|  | TBT | 254 |
|  | TCDD | 215 |
|  | Control | 1326 |
| **Tissue** | Blood | 929 |
|  | Brain | 120 |
|  | Cortex | 180 |
|  | Heart | 24 |
|  | Liver | 1769 |
|  | Lung | 47 |
|  | Skin | 22 |
| **Age** | 3 weeks | 719 |
|  | 5 weeks | 35 |
|  | 20 weeks | 1959 |
|  | 40 weeks | 378 |

### Genomic Pipelines and Quality Control

The TaRGET II data portal also provides access to protocol documents for various assays, including standard operating procedures for handling mice and tissue samples, as well as lab-specific library construction protocols for different tissues across assays. It includes details on the data use policy for external users, quality control (QC) parameters, the tools used, and the common file formats generated across assays. Additionally, the respective experiment library construction and operating protocol documents are made available within the portal.

To ensure reproducibility of analysis, the portal hosts Singularity^13^ images for assay-specific data processing pipelines. These images are self-contained, single-file representations of the complete computational environment. They encapsulate all necessary components—including software, libraries, and the operating system— ensuring that the pipeline runs consistently across different computing platforms.

### Data Portal Services

In the upcoming sections, we outline some of the primary functionalities of the data portal. An essential characteristic of an effective data portal is its capacity to present an overview of the types and quantities of available data, helping users swiftly identify relevant datasets. Besides uniformly processed raw data, comprehensive metadata is gathered and curated, facilitating searches across different data aspects.

### Metadata Overview

Each experiment on the data portal is accompanied by standardized, curated metadata. This includes key attributes such as assay type, tissue, age, sex, mouse strain, and the laboratory responsible for administering the environmental toxicant. An internal mouse ID is also provided to track the source of both target and surrogate tissues. Additionally, the metadata captures detailed treatment information, including the specific toxicant used, exposure dosage, and the exposure paradigm—that is, how the toxicant was administered and the mechanism by which it was delivered.

### Browsing Datasets

The portal’s landing page is designed for intuitive navigation and streamlined access to experimental data. In addition to the top navigation bar, the interactive bar and pie charts enable users to explore datasets by assay type, tissue type, and exposure. Clickable dataset matrices located below these visualizations offer an additional navigation layer, allowing users to browse experiments across two metadata dimensions. Each matrix cell displays the number of experiments corresponding to its row and column definitions. Clicking a cell directs users to the **Explore Page**, with filters pre-applied to display only the relevant experiments for detailed review. Similar dataset matrices are available for Illumina BeadChip arrays^12^ and for ChIP-seq datasets targeting both active and repressive histone marks. For full access to all available data, users can click “Show All Data” to browse the complete set of experiments, as shown in **Figure 1**.

**Figure 1:**
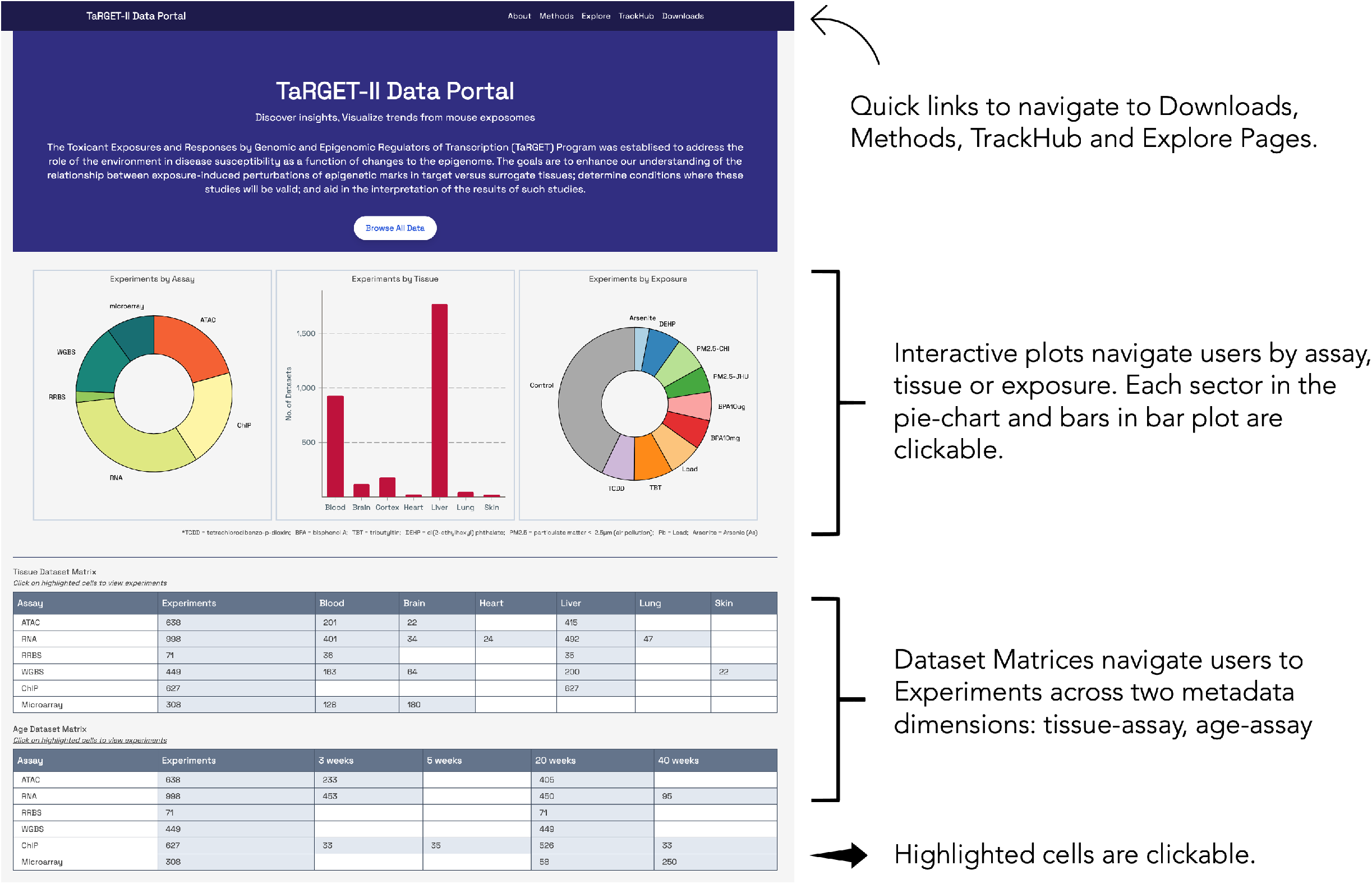
The TaRGET II Data Portal landing page. Interactive plots and dataset matrices provide an easy navigation to experiments of interest. Highlighted cells in the matrix list the number of experiments available at those two-dimensional row and column definitions. Users can click on “Browse All Data” or click on “Explore” in the top navigation bar to view the complete set of experiments.

Within the **Explore Page**, datasets can be further refined using additional filters or by conducting a keyword-based search. The search bar, located in the upper-left corner, supports full-text queries powered by Elasticsearch, offering real-time autocomplete suggestions as users type. Directly below the search box, standard multi-modality filters allow users to narrow datasets by exposure, assay, tissue, age, sex, and originating lab. The displayed results, in the right panel, update automatically in response to filter selections and search inputs. Each row in the results table represents a clickable experiment entry, providing a metadata preview and linking to the corresponding experiment detail page, as shown in **Figure 2**.

**Figure 2:**
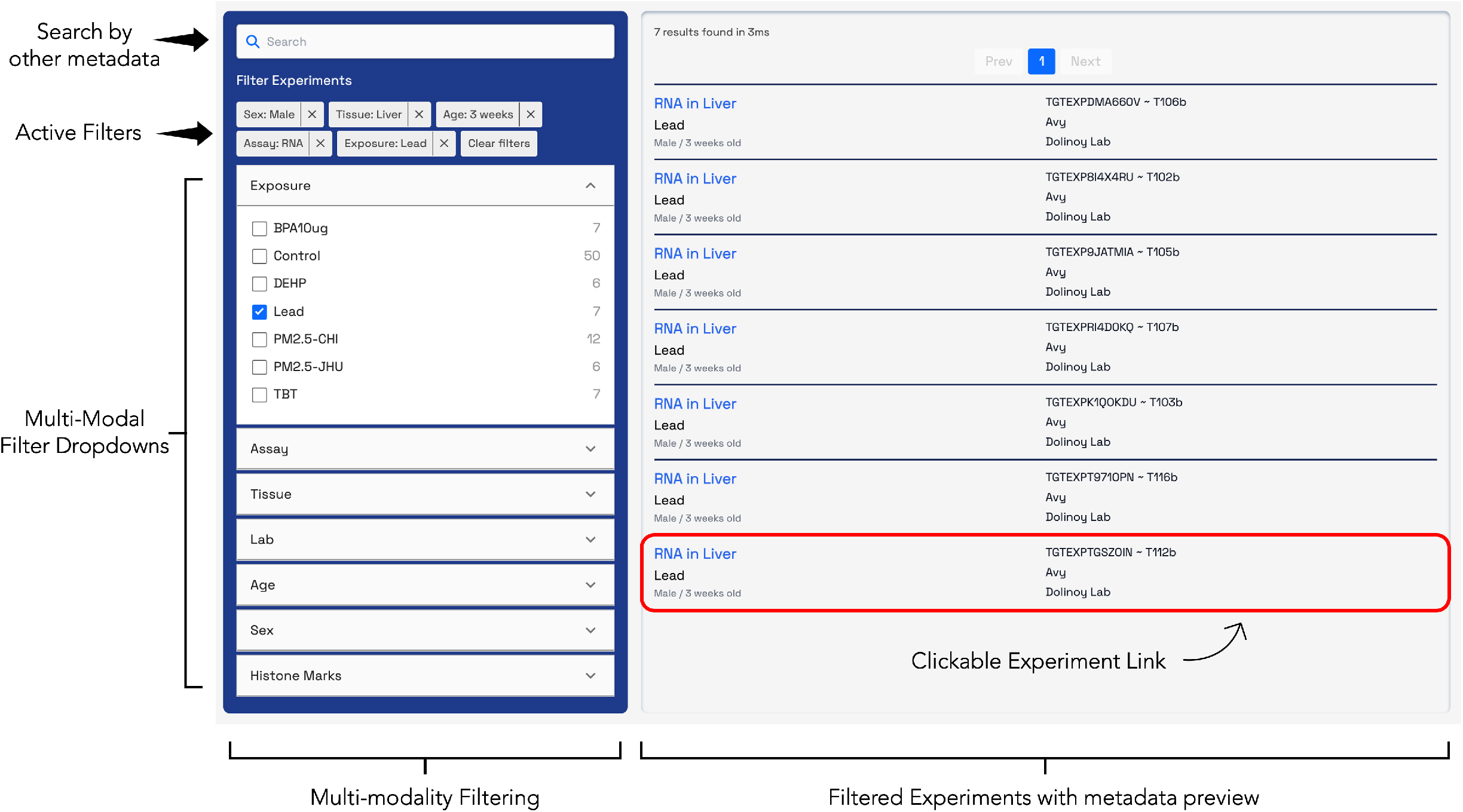
Filters are applied for male mice from the Dolinoy lab, focusing on livers exposed to Lead that were extracted at 3 weeks, with RNA profiling. This search yielded seven experiments listed in the table on the right, accompanied by a preview of relevant metadata, including the selected modalities and experiment IDs.

### Experiment pages

Clicking on any experiment link in the results table will direct users to the experiment page, where more extensive metadata is available. If the dataset has been published, a reference publication will be featured at the top.

Further down the page, several sections provide detailed information about different aspects of the experiment, typically including Processed Files, Raw Files, Attributions, Details, and Warnings or Commendations. Depending on the assay, sections may also include sequencing statistics, tools, tool versions, and the commands used for data processing. Standard quality control metrics and plots are displayed following the sequencing statistics section, as shown in **Figure 3**.

**Figure 3:**
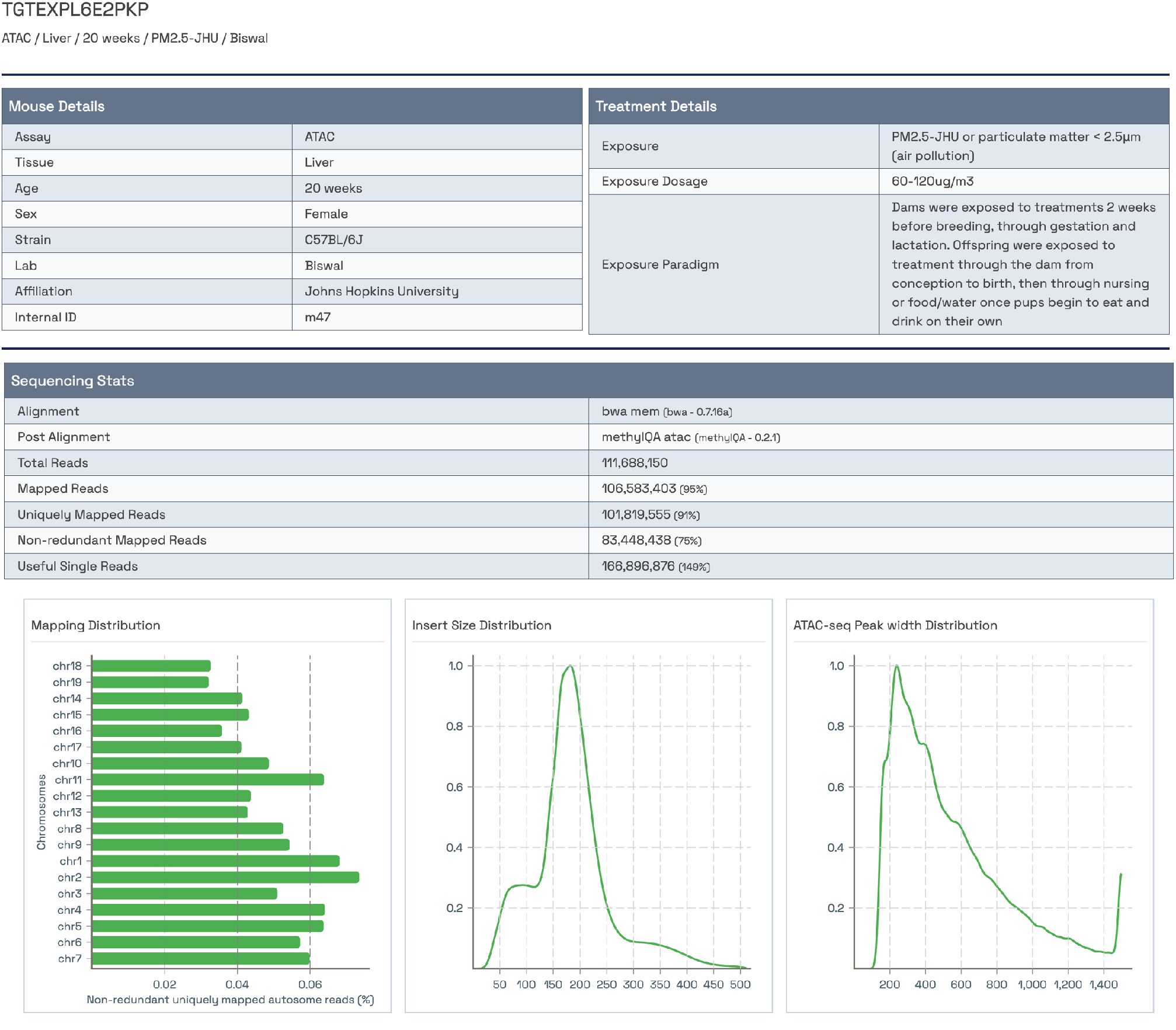
Experiment page section showing attributions and treatment details, followed by sequencing statistics and assay-specific quality control plots.

At the bottom of the page, various file types are available for download through “Direct Download” links and a “Bulk Download” option (as shown in **Figure 4**). The cart icon appears in the navigation bar, displaying the number of datasets added. Clicking the cart icon lists the datasets and allows users to remove individual datasets or completely empty the cart. It also provides instructions for downloading the datasets in the cart. Both raw and processed data, along with common analysis result files, can be downloaded. If applicable, identifiers to other databases that may host the data, such as GEO^14^, are also included. Furthermore, the Downloads page provides alternate options to access all metadata and processed files.

**Figure 4:**
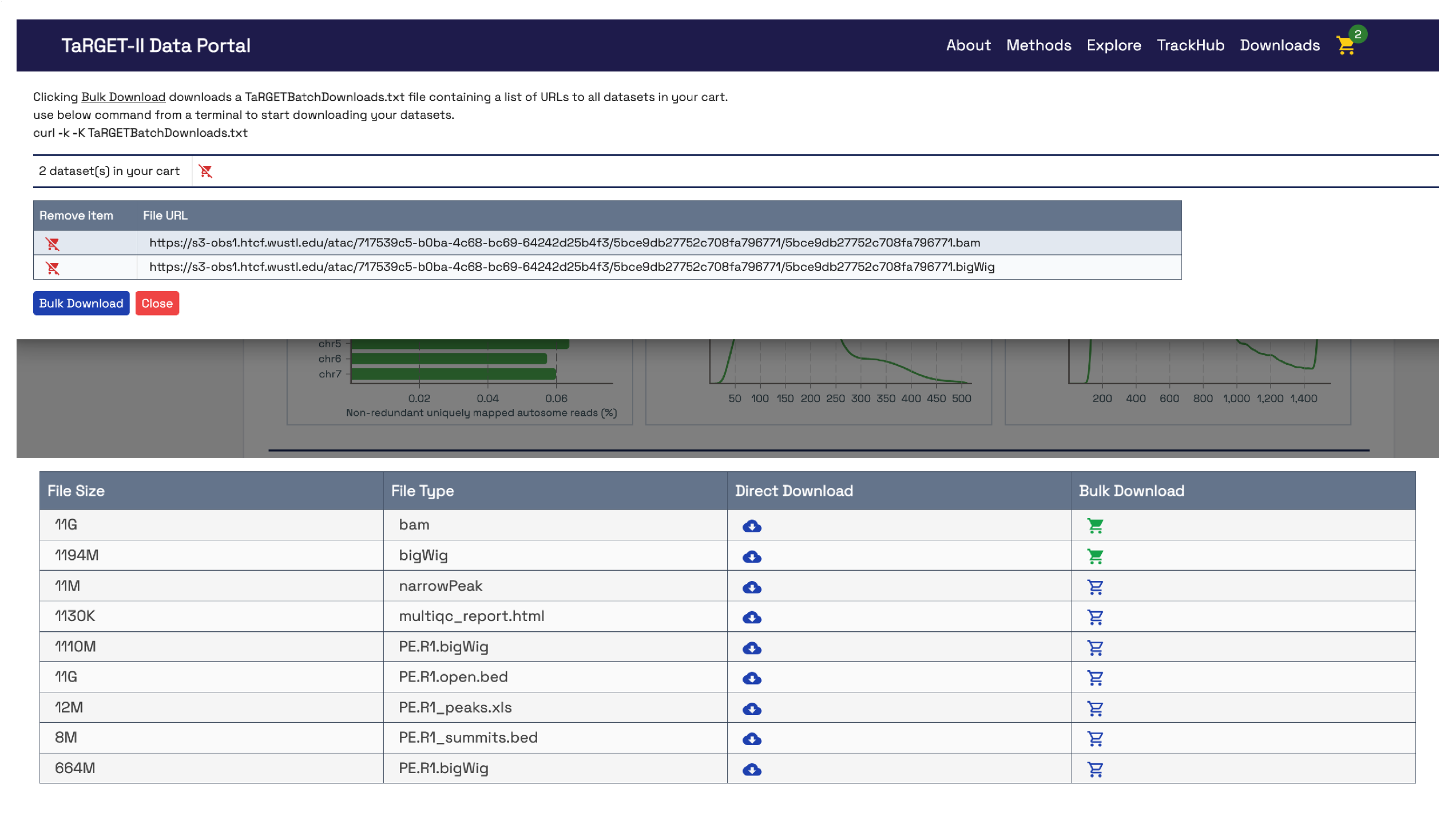
The Experiment page allows users to directly download data files or add them to a download cart. Items in the cart are shown in the navbar. Clicking the cart reveals the datasets and download instructions for bulk downloading all cart items.

### Visualization on the WashU Epigenome Browser

The WashU Epigenome Browser is specifically designed for visualizing, exploring, and generating hypotheses from multi-omics data. In addition to various preconfigured annotation tracks, assay-specific tracks are available through our TaRGET II track hub, enabling visualization of chromatin accessibility, DNA methylation, and gene expression data^15^.

Additionally, a “TrackHub” button at the top directs users to the WashU Epigenome Browser website, linked to the TaRGET II track hub, where they can select multiple tracks for simultaneous visualization. A short tutorial demonstrating how to navigate the WashU Epigenome Browser interface is also available at https://www.youtube.com/watch?v=FzzUT7YEqmA. The close integration of the WashU Epigenome Browser within the portal allows users to utilize the portal’s search capabilities to identify files of interest for visualization, as shown in **Figure 5**.

**Figure 5:**
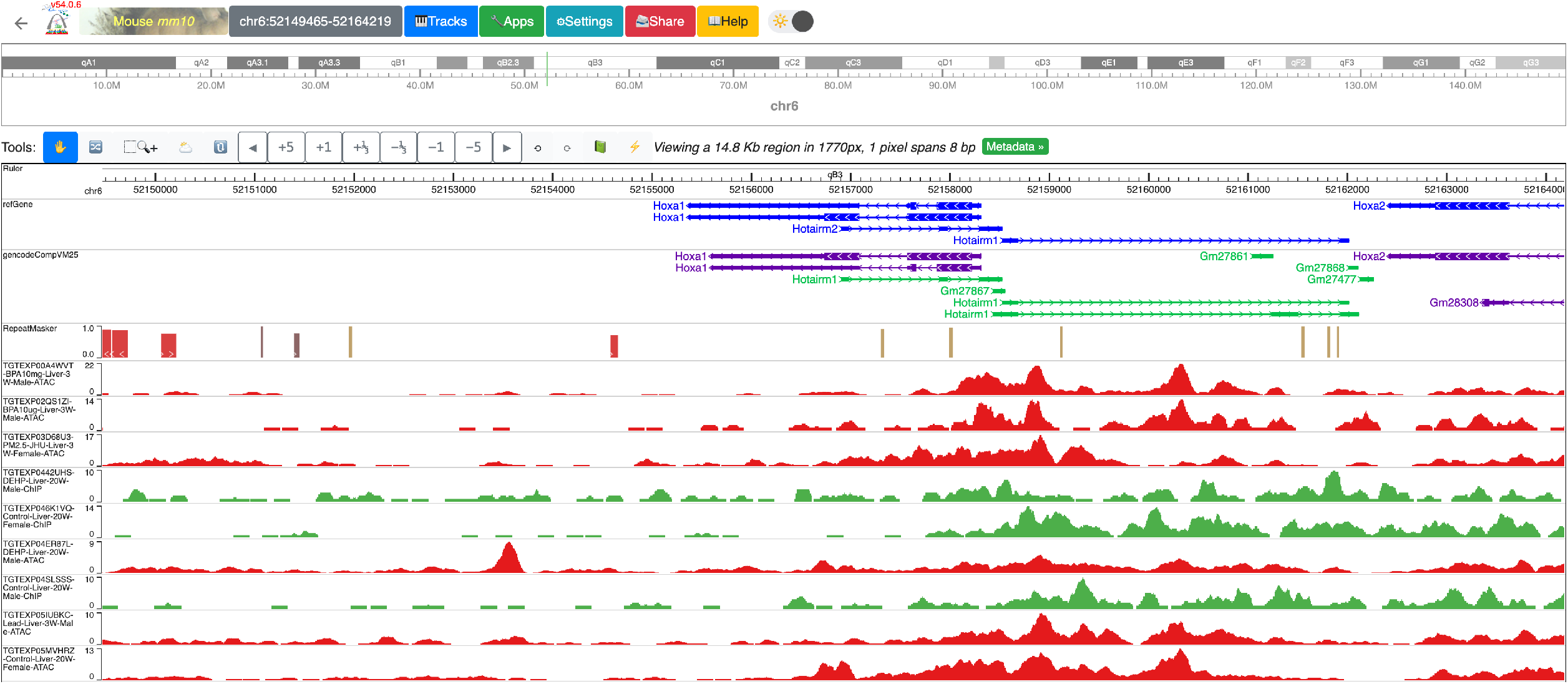
TaRGET II track hub on the WashU Epigenome Browser. The top bar of the browser lists the mouse genome build (mm10) being used, followed by the genomic region, followed by extensive options for loading and manipulating tracks. NCBI’s RefSeq and gencode gene annotation tracks, along with RepeatMasker tracks are made available by default. In this example, the green tracks represent ChIP-seq data, and red tracks represent ATAC-seq data across multiple experiments. The track labels are to the left, displaying the experimentID, exposure, tissue, age, sex and assay.

### Portal Architecture

The TaRGET II consortium was established to generate extensive and diverse exposomes. Recognizing the diverse backgrounds of scientists, the consortium prioritized making data easily discoverable and ensuring that datasets across different modalities are accessible to everyone in the research community. This vision required us to develop a robust and flexible architecture that is modular, responsive, and capable of scaling to accommodate future needs.

The software architecture of the TaRGET II data portal is entirely cloud-based, reflecting the trend of performing data analysis in the cloud rather than downloading large datasets to local servers. This approach facilitates the integration of analytical tools within the cloud environment where the data resides. The infrastructure design adheres to FAIR data standards^16^ and draws inspiration from established data coordination centers (DCC) and data models^17,18^, such as ENCODE’s snoVault ^19,20^ and 4DN’s fourfront^21^. For data storage, the portal primarily utilizes AWS S3, where each quality control pipeline generates an experiment-specific JSON file, along with processed files for download and visualization.

The data portal is developed using the React framework, with the compiled website hosted on AWS CloudFront. All metadata is indexed using AWS Elasticsearch, enhancing search capabilities and accessibility. This robust architecture ensures that the TaRGET II data portal meets the needs of the scientific community by providing a user-friendly interface for accessing and analyzing complex datasets efficiently.

Metadata is submitted by users through standardized spreadsheet forms, while associated data files are uploaded to cloud storage. Once both metadata and files are in place, automated processing pipelines—packaged as Singularity containers—can be executed either on AWS or on on-premises high-performance computing (HPC) systems. All quality control (QC) results and associated metadata are converted to JSON format and indexed using Elasticsearch. The data portal accesses this indexed content via an API Gateway, enabling efficient retrieval and display. External researchers can explore and access the data directly through the portal’s user interface. Download options are available through GEO, AWS Open Data, and directly from the portal itself (as shown in **Figure 6**).

**Figure 6:**
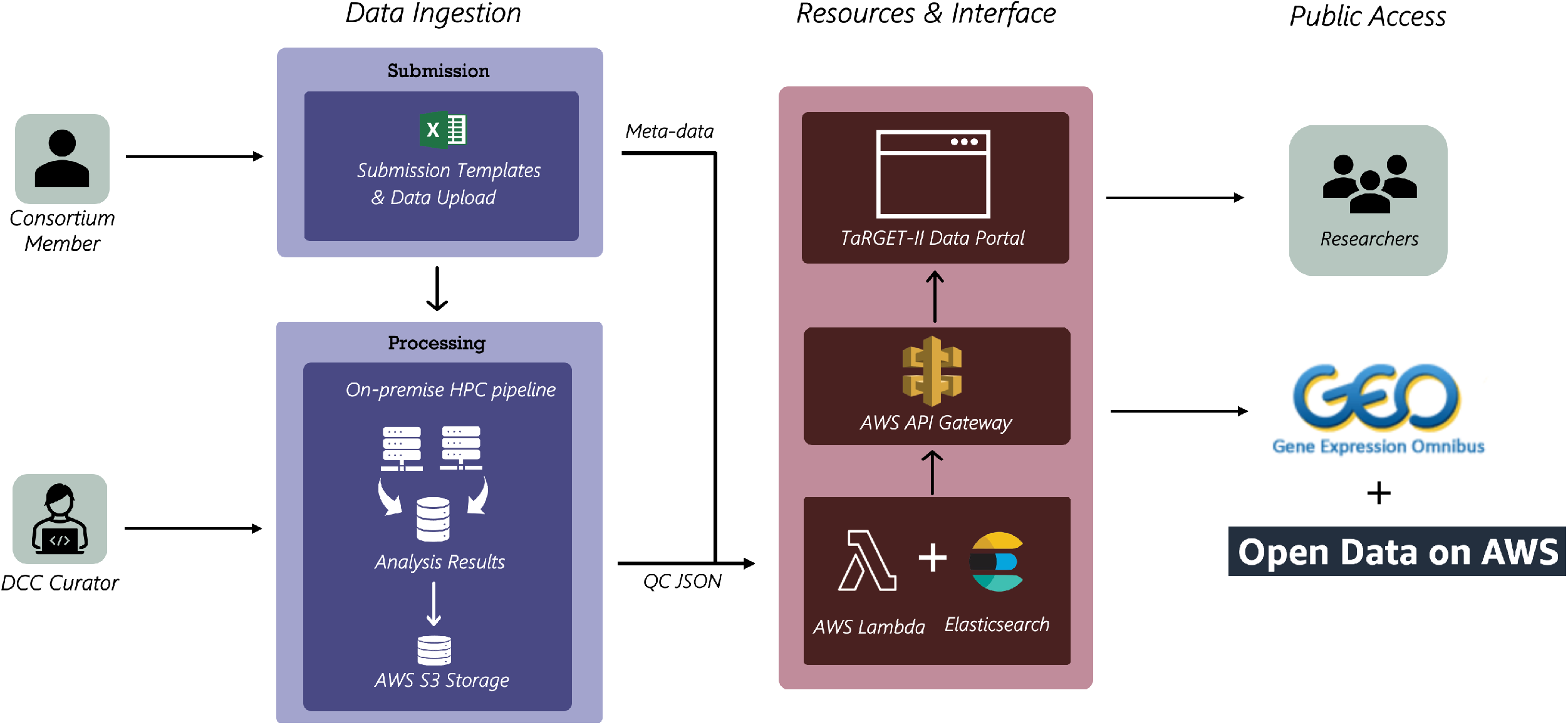
TaRGET II Data Portal architecture. During the Data Ingestion stage, consortium members upload datasets to the cloud along with corresponding metadata forms. A DCC curator then reviews these submissions and initiates the appropriate analysis pipeline. In the Resources and Interface stage, the resulting analysis outputs and metadata are transformed into structured JSON format for indexing and deployment on the public portal. Once this process is complete, the experiments become publicly accessible without the need for login or account creation.

## Discussion

The TaRGET II Data Portal serves as a comprehensive resource for studying the effects of environmental toxicant exposure on the epigenome and disease susceptibility. Designed to accommodate multi-omics datasets, the portal utilizes a cloud-compliant technology stack that ensures scalability for both current and future consortium use cases. It offers users the ability to browse or search for datasets efficiently, with features such as a full-text search powered by Elasticsearch and multi-modality filters for exposure, assay, tissue, age, sex, and lab of origin. This streamlined search functionality enhances user experience and allows for quick access to relevant experiments.

Once users locate datasets, they can click on specific results to access detailed experiment pages containing extensive metadata. These pages provide essential information, including processed and raw files, quality control metrics, and relevant guidelines for data usage, particularly for unpublished datasets. Users can download both raw and processed data files directly or opt for bulk downloads using the cart feature. The portal also includes identifiers for external databases, such as GEO, to further facilitate data accessibility and cross-referencing.

To enhance data visualization, the portal integrates with the WashU Epigenome Browser, allowing users to explore multi-omics data effectively. The “TrackHub” button provides access to the WashU Epigenome Browser website for more comprehensive visualization options. A tutorial playlist is available to assist users in navigating the browser interface. This integration enables researchers to leverage the portal’s search capabilities to identify and visualize data of interest, fostering hypothesis generation and exploration in the context of environmental health research.

## Data Availability

TaRGET II data is part of the AWS Open Data Sponsorship Program and contains data sets that are publicly available for anyone to access and use at: https://registry.opendata.aws/targetepigenomics/.

Individual data files can also be downloaded from the Experiment page within the TaRGET II Data portal using the direct download link or by adding multiple files to a download cart. All the metadata related to all experiments is also available as a comma-separated file (.csv) within the portal’s download page: https://data.targetepigenomics.org/downloads. For unpublished datasets, they remain available for use under the following data usage guidelines: (1) contact the data-generating lab to discuss potential coordinated publication, (2) cite the TaRGET white paper ^7^and portal, and (3) acknowledge the lab that generated the data. A comprehensive data use policy can be found in the “methods” tab on the data portal.

Tutorials and documentation on how to use the TaRGET II data portal are available in the supplemental materials.

## Code Availability

The TaRGET II data portal software is publicly available at https://github.com/twlab/target-portal, under the MIT license. Setup and deployment instructions are made available within the README of the git repository.

## Notes

### Competing Interest Statement

The authors have declared no competing interest.

### Funding

- National Institute Health: U24ES026699

