## supplementary file for "A comprehensive data repository of environmental toxicants exposed mouse epigenomes from the TaRGET II Consortium"

### Supplemental Tutorial:

### Walk-through on using the TaRGET-II Data Portal.

This tutorial has 7 steps demonstrating basic navigation, applying filters, exploring available attributions, downloading datasets via the download cart.

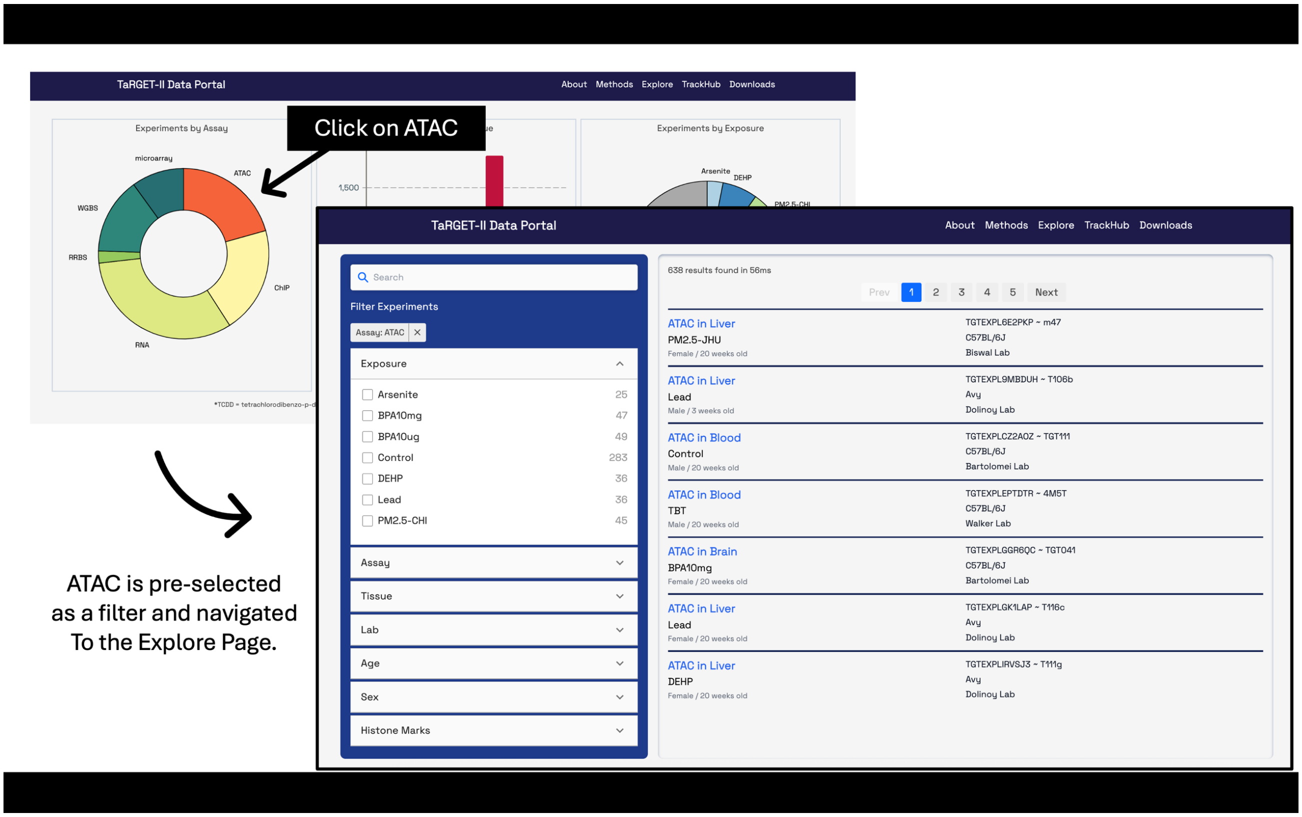


**Step-1:** On the landing page, lets click on the "ATAC" region of the Pie chart. Upon clicking, the users are navigated to the explore page with the filter preselected.

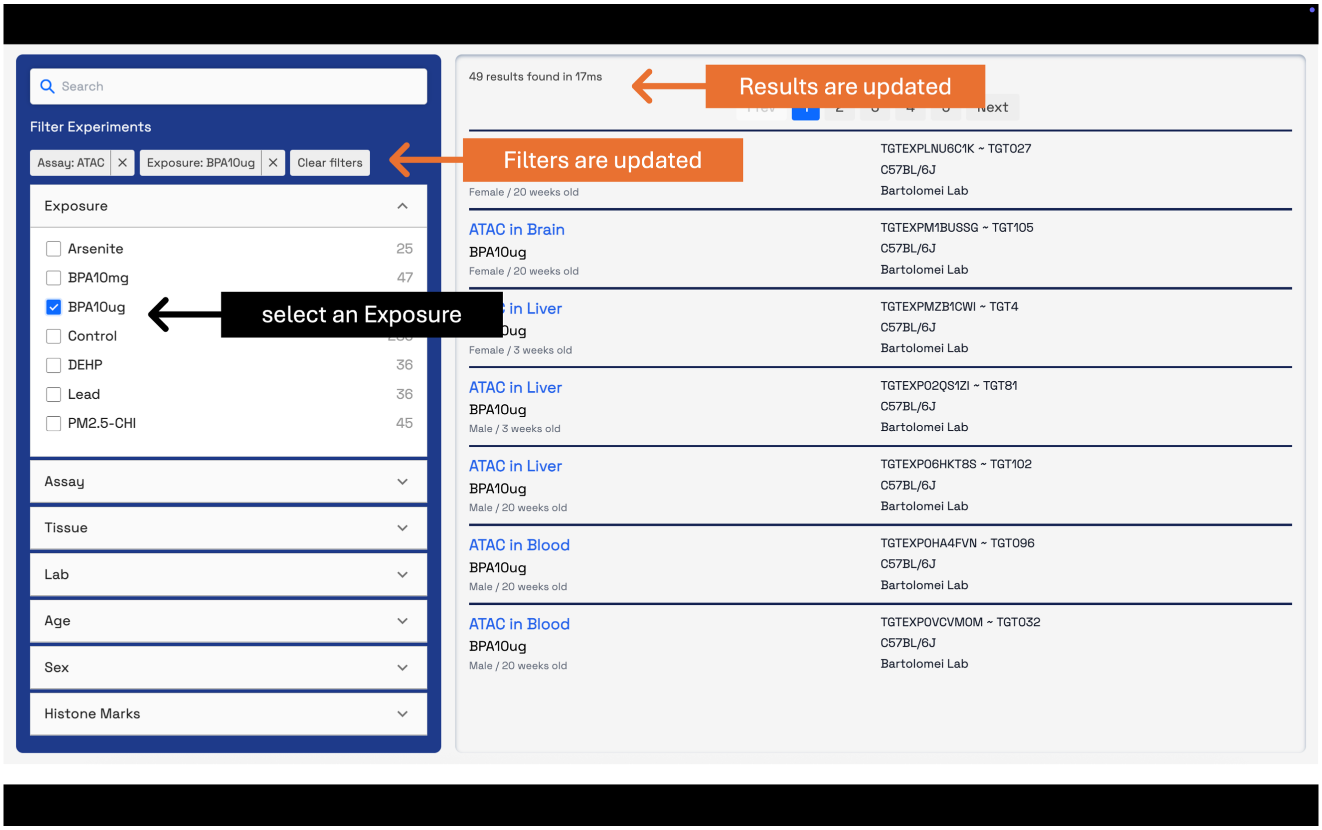


**Step-2:** Now let’s select an exposure “BPA10ug”, this updates the active filters, and the results also change based on these active filters.

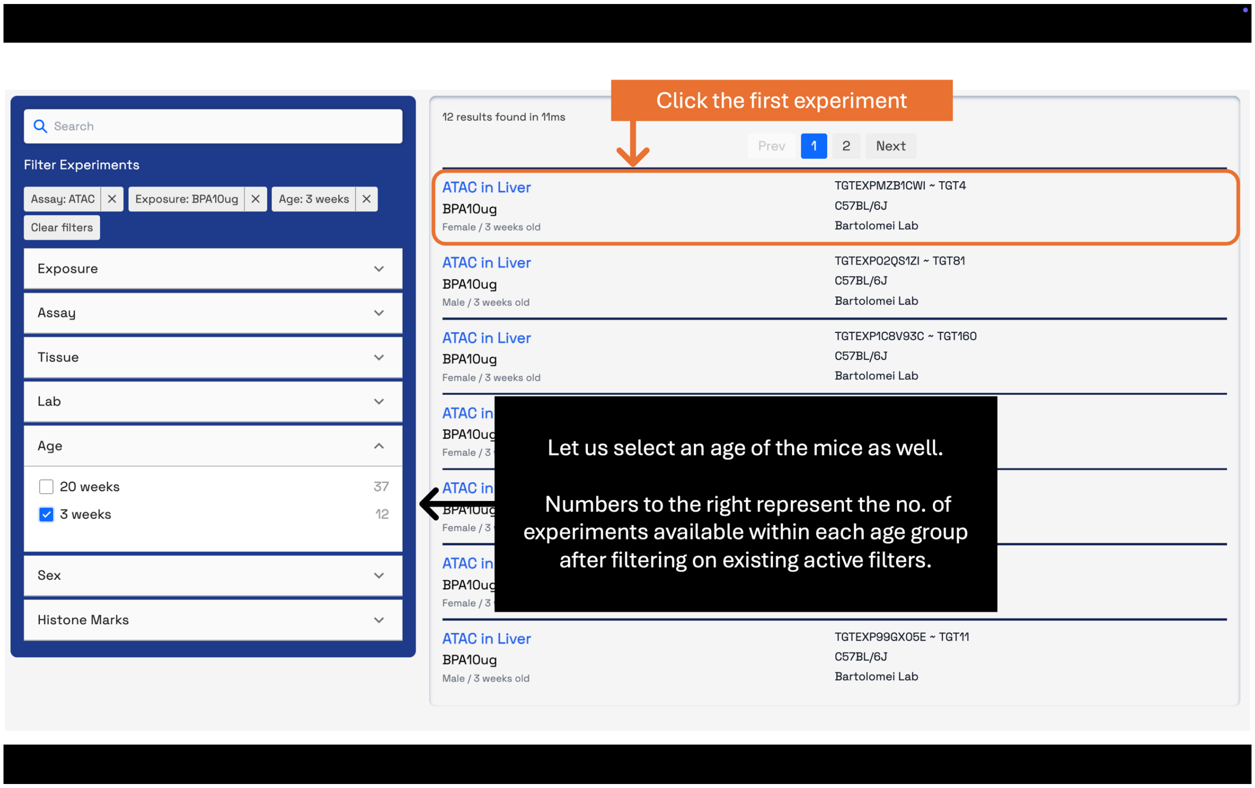


**Step-3:** Now under "Age" filters we can see that based on our current filters there are only 2 age time points available. 3 weeks and 20 weeks. Let's select 3 weeks. The numbers on the right for each age time point, represent total no. of experiments available in that category. Now, click on the first experiment to get to that experiment's "Experiment Page".

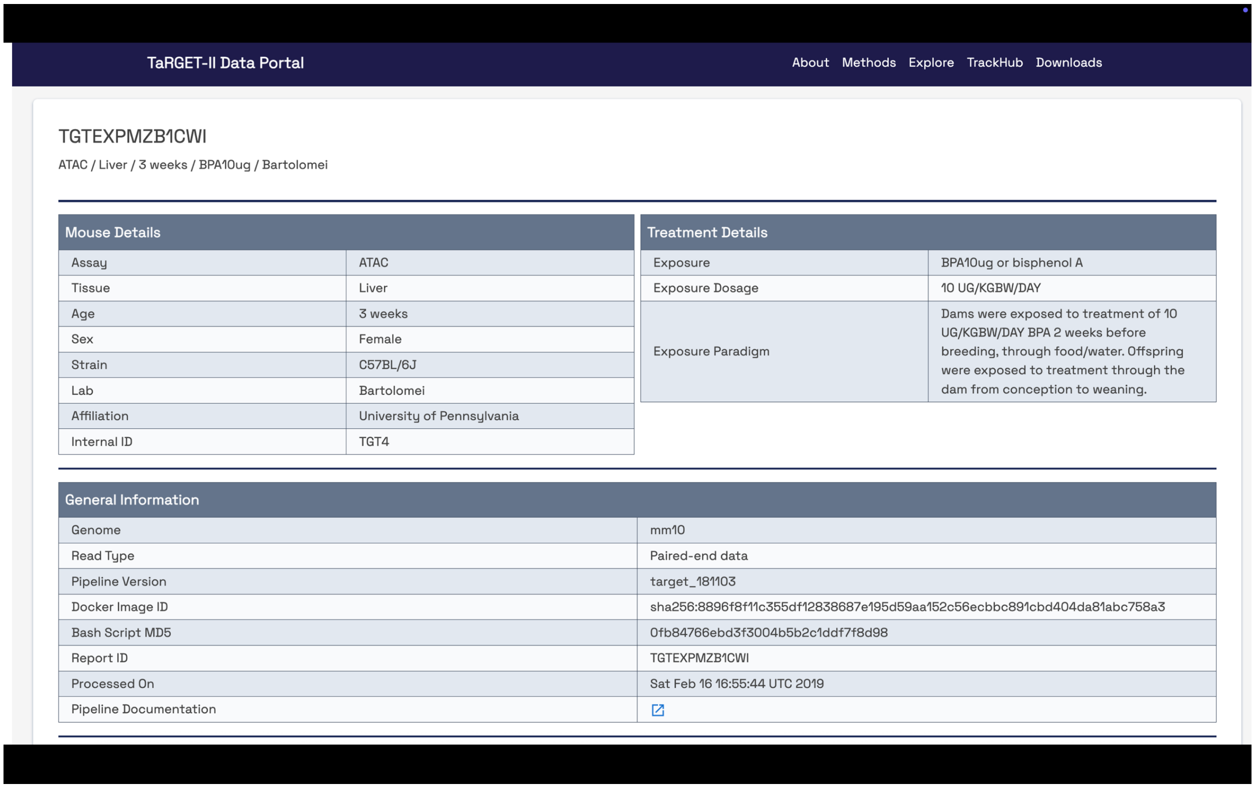

TaRGET-II Data Portal — About | Methods | Explore | TrackHub | Downloads

TGTEXPMZB1CWI

ATAC / Liver / 3 weeks / BPA10ug / Bartolomei

| Mouse Details | |
|---|---|
| Assay | ATAC |
| Tissue | Liver |
| Age | 3 weeks |
| Sex | Female |
| Strain | C57BL/6J |
| Lab | Bartolomei |
| Affiliation | University of Pennsylvania |
| Internal ID | TGT4 |

| Treatment Details | |
|---|---|
| Exposure | BPA10ug or bisphenol A |
| Exposure Dosage | 10 UG/KGBW/DAY |
| Exposure Paradigm | Dams were exposed to treatment of 10 UG/KGBW/DAY BPA 2 weeks before breeding, through food/water. Offspring were exposed to treatment through the dam from conception to weaning. |

| General Information | |
|---|---|
| Genome | mm10 |
| Read Type | Paired-end data |
| Pipeline Version | target_181103 |
| Docker Image ID | sha256:8896f8f11c355df12838687e195d59aa152c56ecbbc891cbd404da81abc758a3 |
| Bash Script MD5 | 0fb84766ebd3f3004b5b2c1ddf7f8d98 |
| Report ID | TGTEXPMZB1CWI |
| Processed On | Sat Feb 16 16:55:44 UTC 2019 |
| Pipeline Documentation | |

**Step-4:** Scroll all the way to the bottom to see available processed files to download.

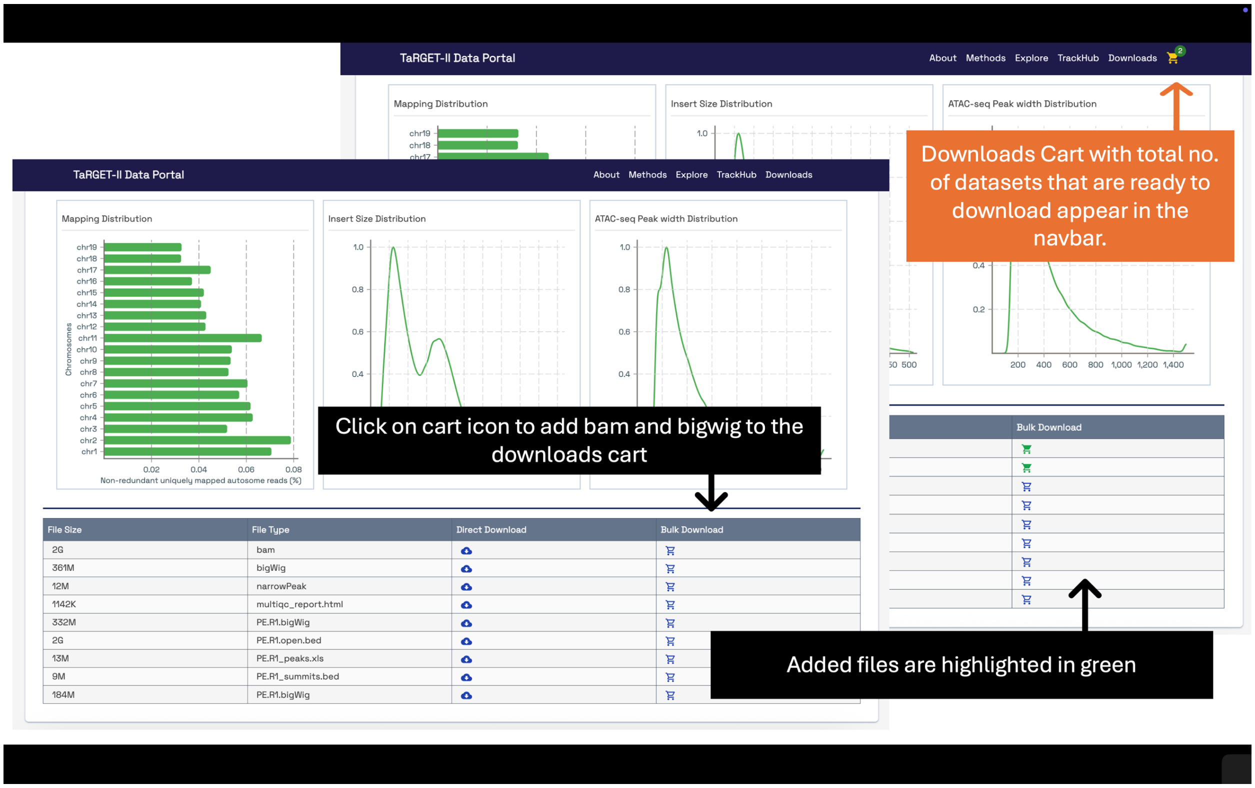


**Step-5:** Clicking on the cart icon for the corresponding file type adds the associated file to the "downloads cart", which appears on the navbar as a "yellow cart" icon. It also changes the cart icon color to green in the processed files table, indicating the file is successfully added to download.

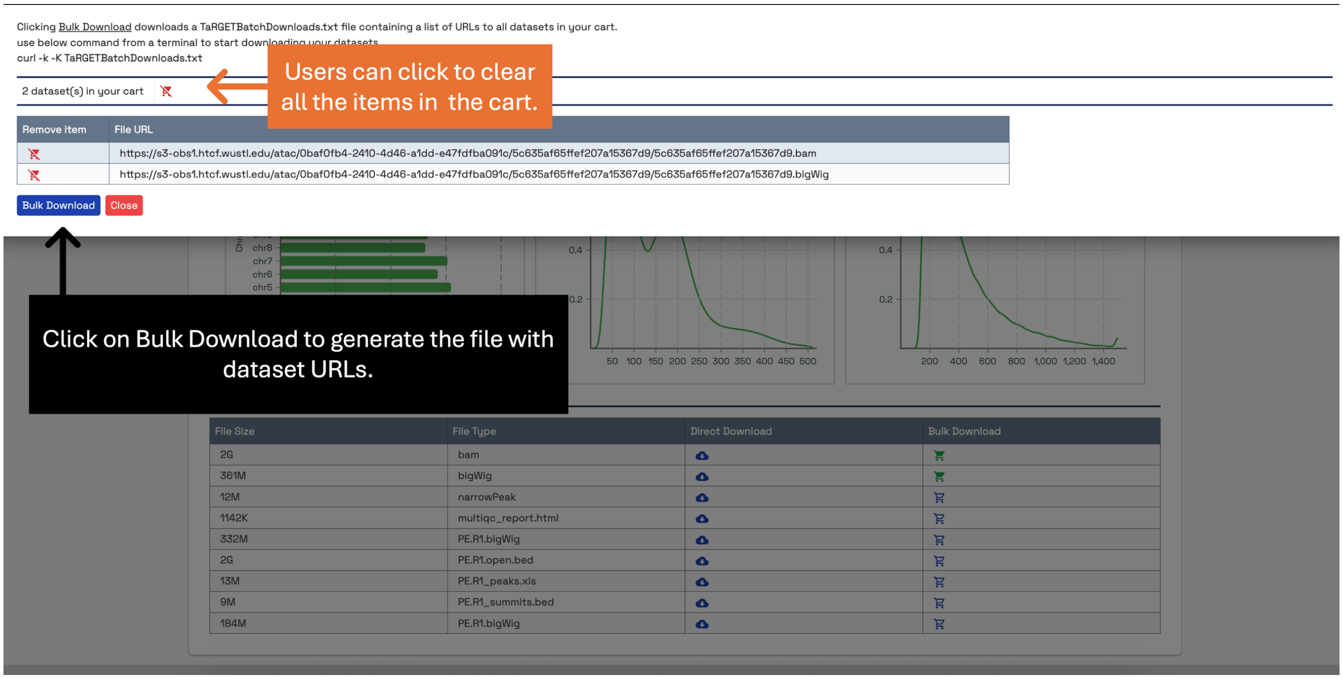


**Step-6:** Clicking on the "yellow cart" icon, shows a dropdown with instructions. Clicking "Bulk Download" will generate a file called "TaRGETBatchDownloads.txt" containing dataset URLs, this file will be automatically saved to your computer in your "Downloads folder". You can decide to remove one or more datasets by clicking the "red cart" icon beside each dataset or clear your cart by clicking the "red cart" icon at the top of the table.

```
> curl -k -K TaRGETBatchDownloads.txt
  % Total    % Received % Xferd  Average Speed   Time    Time     Time  Current
                                 Dload  Upload   Total   Spent    Left  Speed
 14 2677M   14  388M    0     0  34.4M      0  0:01:17  0:00:11  0:01:06 33.8M
```

**Step-7:** Open a terminal application on your computer, navigate to the folder containing the "TaRGETBatchDownloads.txt" file and run the provided curl command as shown. You should start downloading the files one at a time as shown above. This might take a while depending on your internet speed and the file sizes of the datasets you are downloading.
